# Single-cell RNA sequencing deconvolutes the *in vivo* heterogeneity of human bone marrow-derived mesenchymal stem cells

**DOI:** 10.1101/2020.04.06.027904

**Authors:** Zun Wang, Xiaohua Li, Junxiao Yang, Yun Gong, Huixi Zhang, Xiang Qiu, Ying Liu, Cui Zhou, Yu Chen, Jonathan Greenbaum, Liang Cheng, Yihe Hu, Jie Xie, Xucheng Yang, Yusheng Li, Martin R. Schiller, Lijun Tan, Si-Yuan Tang, Hui Shen, Hong-Mei Xiao, Hong-Wen Deng

**Author notes:** Corresponding author Hong-Wen Deng, Ph.D., Professor, Edward G. Schlieder Endowed Chair, Director, Tulane Center for Bioinformatics and Genomics, Department of Biostatistics and Bioinformatics, School of Public Health and Tropical Medicine, Tulane University, 1440 Canal St., Suite 1610, New Orleans, LA 70112, United States of America, u, Hong-Mei Xiao, M.D., Ph.D., Human Genetics and Reproductive Medicine, School of Basic Medical Science, Central South University (China), 173 Tongzipo Rd., Changsha, 70118, China. These authors contributed equally to this work (Zun Wang, Xiaohua Li).

## Abstract

Bone marrow-derived mesenchymal stem cells (BM-MSCs) are multipotent stromal cells, which have a critical role in the maintenance of skeletal tissues such as bone, cartilage, and the fat found in bone marrow. In addition to providing microenvironmental support for hematopoietic processes, BM-MSCs can differentiate into various mesodermal lineages including osteoblast/osteocyte, chondrocyte, and adipocyte cells that are crucial for bone metabolism. While BM-MSCs have high cell-to-cell heterogeneity in gene expression, the cell subtypes that contribute to this heterogeneity *in vivo* in humans have not been characterized. To investigate the transcriptional diversity of BM-MSCs, we applied single-cell RNA sequencing (scRNA-seq) on freshly isolated CD271^+^ BM-derived mononuclear cells (BM-MNCs) from two human subjects. We successfully identified LEPR^hi^CD45^low^ BM-MSCs within the CD271^+^ BM-MNC population, and further codified the BM-MSCs into distinct subpopulations corresponding to the osteogenic, chondrogenic, and adipogenic differentiation trajectories, as well as terminal-stage quiescent cells. Biological functional annotations of transcriptomes suggest that osteoblast precursors may induce angiogenesis coupled with osteogenesis, and chondrocyte precursors may have the potential to differentiate into myocytes. We discovered transcripts for several cluster of differentiation (CD) markers that were highly expressed (e.g., CD167b, CD91, CD130 and CD118) or absent (e.g., CD74, CD217, CD148 and CD68) in BM-MSCs and could be novel markers for human BM-MSC purification. This study is the first systematic *in vivo* dissection of human BM-MSCs cell subtypes at the single-cell resolution, revealing insight into the extent of their cellular heterogeneity and bone homeostasis.

## Introduction

The human bone tissue is a complex system that consists of diverse cell types including osteoblast/osteocyte, osteoclast, and chondrocyte (collectively known as “bone cells”), together with various supporting cells such as adipocyte, fibroblast, and hematopoietic cells among others. A delicate balance of bone formation/resorption is critical for maintaining bone health, and therefore bone cells must work together to maintain bone strength and mineral homeostasis. Despite the extensive study of bone cells, their underlying biology remains poorly understood. While osteoclasts are of hematopoietic origin and derived from the “monocyte/macrophage–preosteoclast–osteoclast” differentiation trajectory^1^, the detailed origins of osteoblast/osteocyte and chondrocyte are not as well characterized.

Currently, the cells that give rise to osteoblast/osteocyte, chondrocyte, and adipocyte are generally referred to as “mesenchymal stromal/stem cells” (MSCs), which are non-hematopoietic bone marrow stromal cells with fibroblast colony-forming unit (CFU-F) and multi-differentiation capacity^2^. Typically, the human bone-marrow derived MSCs (BM-MSCs) are isolated with a combination of non-specific cell-surface markers such as high expression of CD271, CD44, CD105, CD73, CD90, and low expression/absent of CD45, CD34, CD14 or CD11b, CD79a or CD19, and human leukocyte antigen HLA-DR^3,4^. Among these markers, CD271 shows great efficiency to sort MSCs either alone or in combination with negative selection of markers such as CD45^5,6^. Additionally, LEPR (leptin receptor, or CD295) is used for isolating BM-MSCs in transgenic labeling mice^7,8^.

Although these cell markers are candidates for isolating BM-MSCs, recent evidence suggests that the BM-MSCs are a heterogeneous group of cells for some cell markers. For instance, Akiyama et al.^9^ demonstrated that a small portion of BM-MSCs express CD45 and CD34, which are traditionally regarded as negative markers. Meanwhile, some studies also suggested that only around 50% of MSCs are positive for CD105^10,11^, a cell marker previously considered universally expressed by MSCs derived from different tissue^12^.

The extent of the cellular heterogeneity among the BM-MSCs is not well-defined, although a few studies have proposed some novel subtypes. One study reported a subset of cultured mouse BM-MSCs that are distinct from regular BM-MSCs based upon differential attachment to plastic culture dishes, proliferation, and self-renewal patterns^9^. Another study examining cultured human BM-MSCs demonstrated that CD264 marks a subpopulation of aging human BM-MSCs with differential fibroblast colony forming efficiency^13^. Several other efforts have attempted to deconvolute the heterogeneity of BM-MSCs through the identification of the differentiation trajectory associated with a given subpopulation. For example, one study found that effective chondrocyte differentiation could only be induced in human MSCA-1^+^CD56^+^ BM-MSCs, while adipocytes are derived only from MSCA- 1^+^CD56^−^ BM-MSCs *in vitro*^14^. Another study identified “skeletal stem cells” in both mice and humans, which give rise to bone, stroma, and cartilage cells *in vivo* in mice, but not adipocytes or myocytes^15,16^.

Single-cell RNA sequencing (scRNA-seq) has recently emerged as a powerful approach to study cell heterogeneity in complex tissues. scRNA-seq measures transcriptional profiles of many cells at single-cell resolution, which can be clustered to distinguish and classify cell subtypes and infer developmental trajectories, as well as identify novel regulatory mechanisms^17,18^. scRNA-seq technology represents a major advancement beyond conventional bulk RNA-seq transcriptomics approach, which attempts to infer biological mechanisms from average gene expression, weighted by the unknown proportions of unknown cell subtypes, across a heterogeneous cell population. Several studies have applied scRNA-seq to bone marrow stroma cells. However, these studies were on mice^7,19^ or cultured cells from human subjects^20,21^, which are less likely to properly represent the transcriptional profile of human primary BM-MSCs *in vivo*^22,23^.

Our work is the first systematic scRNA-seq analysis of freshly isolated human CD271^+^ bone marrow mononuclear cells (BM-MNCs). We successfully identified LEPR^hi^CD45^low^ BM-MSCs in the CD271^+^ BM-MNC population, revealing distinct subpopulations in LEPR^hi^CD45^low^ BM-MSCs along with their differentiation relationships and functional characteristics. By comparing the expression pattern of LEPR^hi^CD45^low^ BM-MSCs with CD45^hi^ hematopoietic cells, we also propose some novel markers for human BM-MSC purification. The findings provide significant insight into the identities and complexities of human BM-MSCs *in vivo*.

## Results

### Cellular heterogeneity of the human CD271^+^ BM-MNCs

To study the transcriptomic diversity of the BM-MSCs, we applied scRNA-seq on freshly isolated CD271^+^ BM-MNCs from the femoral shafts-derived bone marrow of two human subjects (one with osteoporosis and the other with osteoarthritis) (**Fig. 1a**). Cells were affinity isolated with CD271 conjugated magnetic microbeads (**See methods**), and mRNA libraries were prepared and sequenced with the 10x Genomics Chromium system. After quality filtering (**Extended Data Fig. 1a-c**), we obtained an expression matrix of 14,494 cells where transcripts for the average number of genes detected per cell was 1,363. There was a strong correlation between the overall gene expression profiles of the two subjects (*R* = 0.96, **Extended Data Fig. 1d,e**), therefore, we combined the data from the two subjects for subsequent analyses. The graph-based clustering divided the cells into 15 distinct clusters (clusters A-O), and their differentially expressed genes (DEGs) of each cluster were identified with the Wilcoxon rank-sum test (**Extended Data Fig. 2a**,**b; Supplementary Table 1: Sheet 1**).

**Fig. 1.**
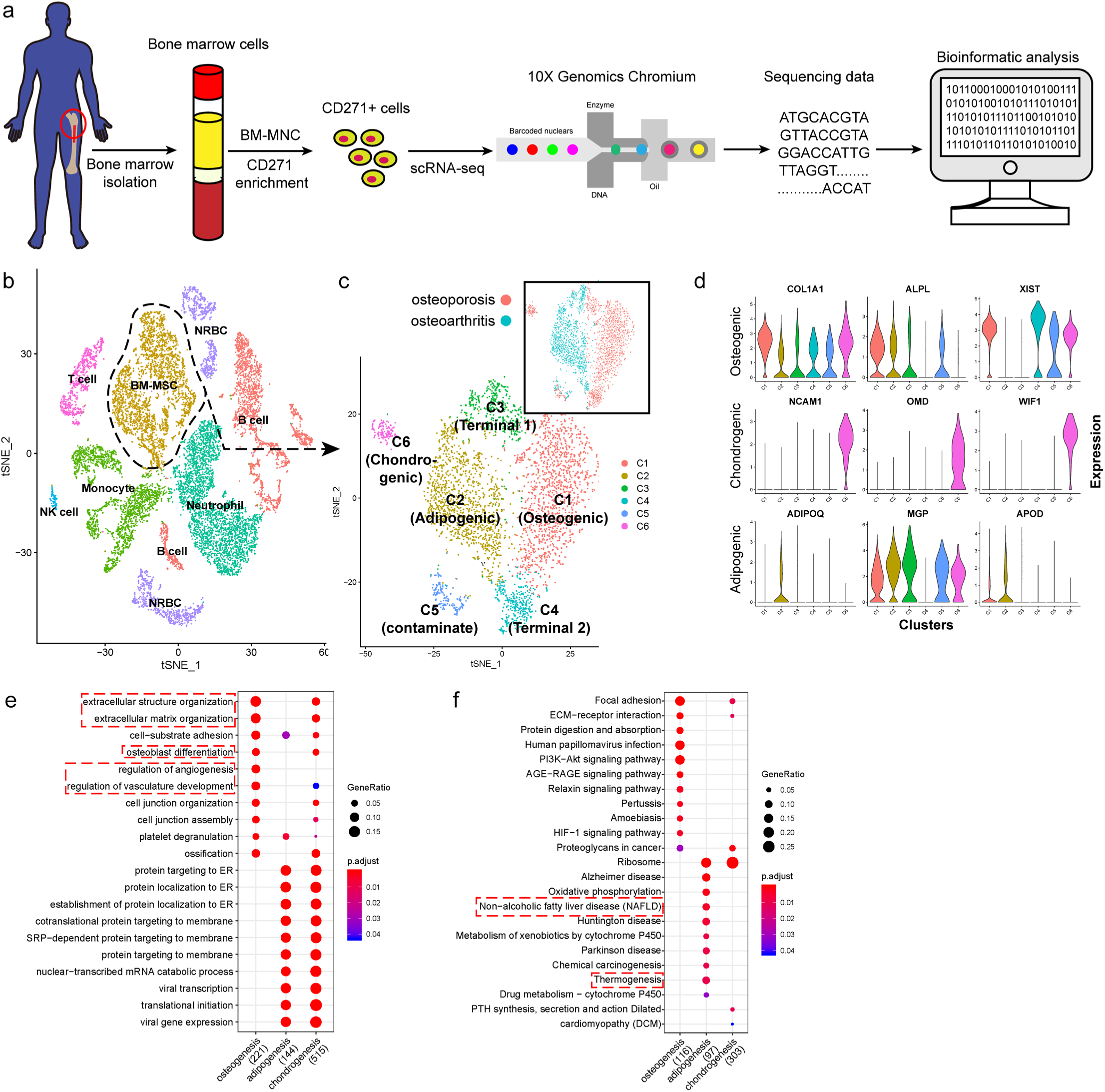
scRNA-seq analysis of the human BM-MSCs. **a**, Schematic summarizing and overview of the study. **b-c**, t-SNE visualization of color-coded clustering of 14,494 human CD271^+^ BM-MNCs. The labeled texts indicate the individual clusters. Dashed lines in (**b**) delineates LEPR^hi^CD45^low^ BM-MSCs, which are further classified into subgroups shown in (**c**). The upper-right t-SNE plot in (**c**) shows the difference in BM-MSCs between the two subjects. **d**, Violin plots showing relative expression levels of selected cluster-specific marker genes for osteoblast (top row), chondrocyte (middle row), and adipocyte (bottom row) precursors, respectively. **e-f**, GO (**e**) and KEGG (**f**) enrichment analyses for osteoblast, chondrocyte, and adipocyte precursors. Dot plot shows the most significant terms. The size of dot indicates the gene ratio (enriched genes / total number of genes). The color indicates the adjusted *p* value for enrichment analysis. Dashed boxes highlight the terms related to MSC functions.

**Fig. 2.**
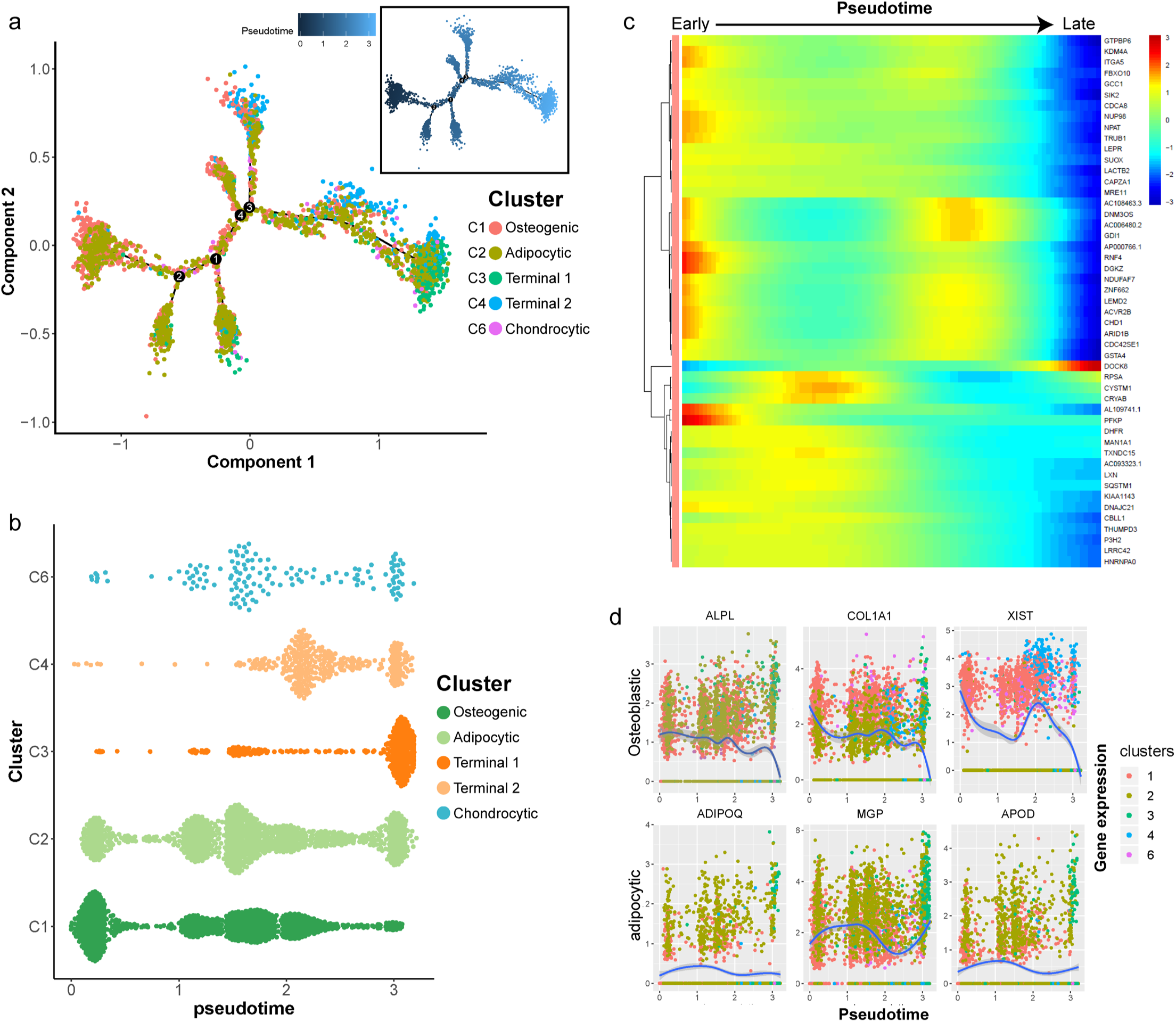
Dynamic Gene Expression Patterns of Human BM-MSCs. **a**, Reconstructed principal component graph of cell differentiation trajectory of BM-MSCs, colored by subpopulation identities. The upper-right trajectory plot in the square indicates the direction of pseudotime. **b**, Distribution of each cell subpopulation along the pseudotime. **c**, Continuum of dynamic gene expression in pseudotime of BM-MSCs. The heatmap shows the top 50 genes with most significant expression changes. Pixel color indicates the expression level (logFC). **d**, Expression level of osteogenic (top) and adipogenic (bottom) genes with respect to pseudotime coordinates. Blue lines depict the LOESS regression fit of the normalized expression values.

Among the cell type clusters, clusters C and D expressed high levels of BM-MSC marker genes, including LEPR (leptin receptor), NGFR (CD271), ENG (CD105), THY1 (CD90), and NT5E (CD73). Notably, LEPR had the strongest expression levels (**Extended Data Fig. 2c**). The remaining clusters are PTPRC (CD45) or HBA1 (hemoglobin-1) positive hematopoietic cells (**Extended Data Fig. 2c**). Specifically, based on the presence of marker: **1)** clusters A and B are CD11b/16/66b^hi^ neutrophils; **2)** clusters F, K, N are CD14^hi^CD16^low/hi^ monocytes; **3)** clusters E, I, L, and M are immunoglobulin^hi^ B cells; **4)** cluster H is CD3^hi^ T cells; **5)** cluster O is CD56^hi^ NK cells; and **6)** clusters G and J are hemoglobin^hi^ nucleated red blood cells (RBCs) (**Fig. 1b; Supplementary Table 1: Sheet 1**). These findings are consistent with previous reports that MSCs are the main source of LEPR expression in human bone marrow and CD271^+^ MNCs also express certain levels of CD45 (**Extended Data Fig. 2c**)^6,24^. By comparing the gene expression pattern between LEPR^hi^CD45^low^ BM-MSCs and other CD45^hi^ hematopoietic cells, we discovered several potential surface markers for isolation of human BM-MSCs such as high expression of CD167b, CD91, CD130, CD118 and low expression or absence of CD74, CD217, CD148, CD68 (**Supplementary Table 1: Sheet 2**). These results demonstrate that CD271^+^ MNCs is a heterogeneous cell population containing many cell types.

### Cellular taxonomy of BM-MSCs

To investigate the cellular heterogeneity within BM-MSCs, we extracted LEPR^+^CD45^-^ cells (clusters C and D, **Extended Data Fig. 2a**) from the original dataset for further analyses. The BM-MSCs were divided into six distinct groups by an unbiased clustering analysis (**Fig. 1c, and Extended Data Fig. 2d**). Based on known cell markers or functional genes, the different subtypes of BM-MSCs were annotated as: **1)** osteoblast precursor (cluster C1, expressing osteogenesis markers including collagen 1 and ALPL^25,26^); **2)** adipocyte precursor (cluster C2, expressing adiponectin and MGP^27,28^); **3)** chondrocyte precursor (cluster C6, expressing CD56 and WIF1^14,29^); and **4)** terminal-stage cells which do not express differentiation markers (clusters C3-C5) (**Fig. 1d**).

We studied the expression and the function of the cluster-specific DEGs of the new BM-MSCs subpopulations (**Supplementary Table 1: Sheet 3**) with interesting results: **1)** besides known markers or functional genes such as ALPL and collagen 1, some novel genes were also highly expressed in the osteoblast precursor cells. For instance, XIST, a long non-coding RNA (lncRNA) that regulates chondrocyte proliferation and apoptosis through MAPK signaling^30^, was highly expressed in the osteoblast precursor, suggesting that XIST might also be a novel regulator for osteogenesis. In addition, MCAM (CD146) was also differentially expressed in osteoblast precursor when compared with other cell subtypes. CD146 was recently regarded as one of the markers for human osteoblast precursor^15^. **2)** With ADPQ (adiponectin) and MGP, APOD (apolipoprotein D) was also highly expressed in the adipocyte precursor. Though APOD is not yet linked with adipogenesis, other members of the apoliproteins, such as APOA and APOE^31,32^ are known to modulate adipocyte metabolism. Therefore, it is conceivable that APOD may also regulate adipogenesis. **3)** Osteomodulin (OMD) was highly expressed in the chondrocyte precursor. Previous reports have shown that OMD induces endochondral ossification through PI3K signaling, regulates the extracellular matrix during bone formation by reorganizing collagen fibrils, and increasing aggrecan expression in chondrocytes^33–35^. Taken together, the findings suggest that OMD may be an important factor regulating chondrogenic differentiation.

To further validate our findings, as well as to study the shared and distinct biological processes between different cell type clusters, we analyzed GO and KEGG for enrichment of DEGs in osteoblast, chondrocyte, and adipocyte precursors (**Supplementary Table 2: Sheets 1-2**). Enrichment terms for bone development in the osteoblastic and chondrocyte precursors were identified including “ossification”, “osteoblast differentiation”, etc. Terms related to adipocytes were enriched in the adipocyte precursors by KEGG pathway enrichment such as “non-alcoholic fatty liver disease” and “thermogenesis” (**Fig. 1e**,**f**)^36,37^. These results demonstrate that human BM-MSCs consist of a heterogeneous cell population with several different subtypes, which are characterized by distinct biological processes and subject phenotype.

In contrast, the remaining subgroups (clusters C3-C5) of the BM-MSCs did not express any differentiation markers, and GO enrichment analyses did not detect any significant terms related to differentiation processes. Ribosomal Protein (RP) gene family, which encodes ribonucleoprotein, were highly expressed in clusters C3 and C4 (**Extended Data Fig. 2d**). Previous evidence suggests that the expression of ribonucleoprotein is required for maintenance of self-renewal of potency stem cells^38^. These clusters were enriched for the GO terms related to ribonucleoprotein, RNA degeneration, and cell apoptosis (**Extended Data Fig. 3a**). These results support the claim that these clusters contain cells at terminal stage and lack the capacity for cellular differentiation. We noted that although cluster C5 had high expression levels of LEPR, a small fraction of the cells in this group also expressed low levels of CD45 and were enriched for immune cell related terms such as “neutrophil cell activation” and “leukocyte migration” (**Extended Data Fig. 2e, and 3a**). This result suggested that CD45^+^ immune cells may have contaminated this cluster. Thus, we excluded this cluster (C5) from further analysis.

**Fig. 3.**
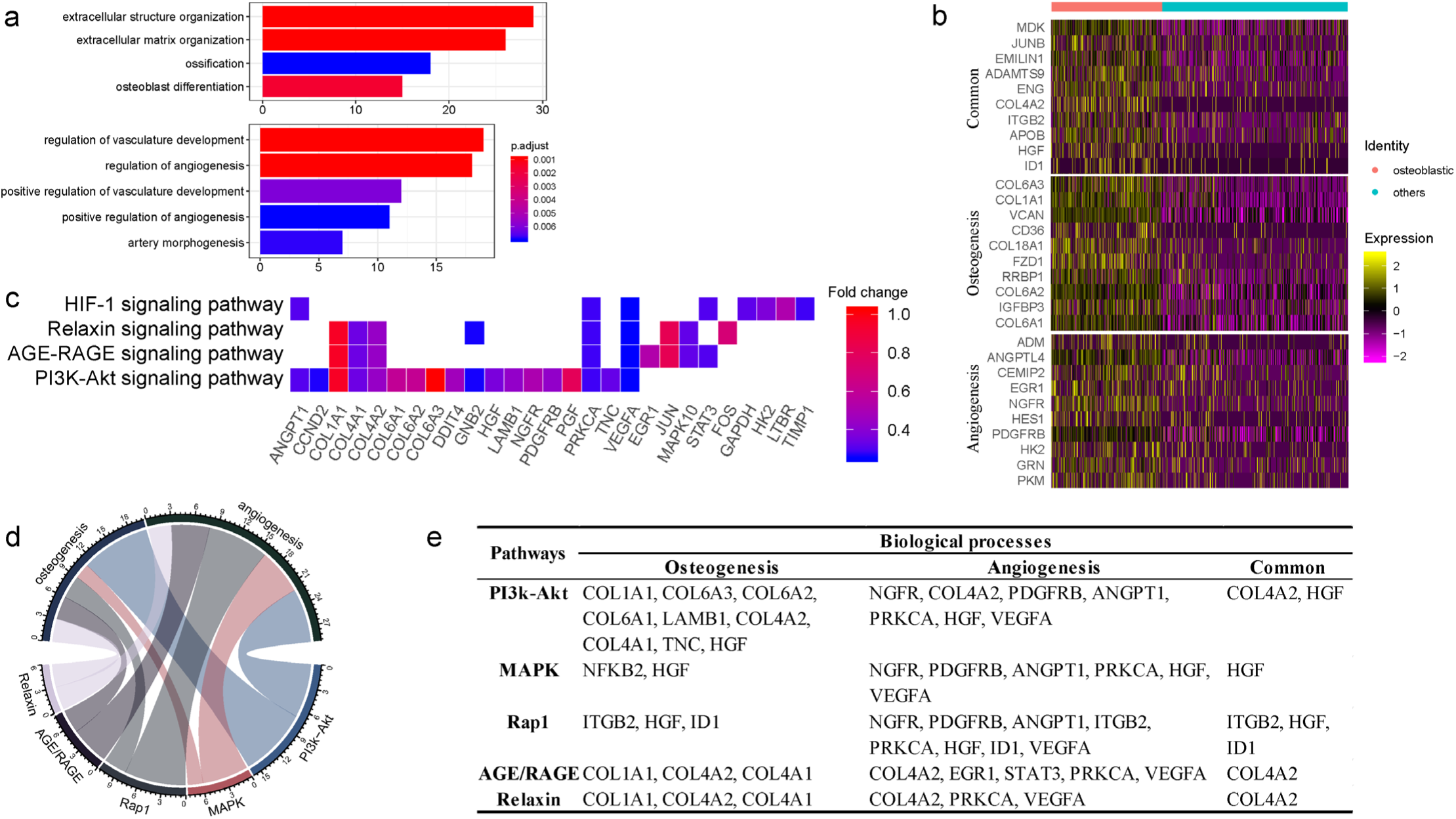
Functional analysis for ALPLhi osteoblast precursor. **a**, Enriched GO terms associated with osteogenesis (top) and angiogenesis (bottom) in osteoblast precursors. Bar chart shows the number of genes enriched in each term. Color indicates the adjust *p* values. **b**, Differential expression of osteogenesis- and (or) angiogenesis-related genes (rows) in osteoblast precursors compared to the other cells. Heatmap shows the top 10 most significant DEGs in each category, where color indicates the relative expression levels between osteoblast precursors and other cells (z-score). **c**, Gene expression pattern in enriched pathways. Squares show enriched DEGs in the corresponding terms (rows). Color indicates the expression value of the DEGs (average logFC). **d-e**, Table of genes in biological processes and pathways. (**d**), Numbers outside the circles indicates the number of genes in that term. Width of curves connecting different terms is proportional to the number of shared genes. (**e**), table of the specific genes enriched in each biological process and pathway.

### Dynamic gene expression patterns at different developmental stages of BM-MSCs

In order to better understand the differentiation relationships between BM-MSCs subtypes, we reconstructed the developmental trajectory by inferring the dynamic gene expression patterns at different developmental stages. The estimated developmental trajectory showed multiple branches, representing multi-lineage differentiation potential of BM-MSCs (**Fig. 2a**). By comparing the distribution of the cell population along the pseudotime, we found that osteoblast precursors (cluster C1) were more enriched in the early stage of pseudotime compared with the other clusters, while adipocyte and chondrocyte cells were evenly distributed along the pseudotime. Meanwhile, clusters C3 and C4 were mostly represented at the later stage of the pseudotime, supporting that these groups contain cells at terminal stages (**Fig. 2b**). Pseudotime ordering of cell type clusters revealed a continuum of gene expression between the early and late stages of BM-MSC differentiation (**Fig. 2c**). When the dynamic gene expression patterns between osteoblast and adipocyte markers are compared, the osteoblast markers decreased over pseudotime, while adipocyte markers remained unchanged or increased (**Fig. 2d**). These findings suggest that osteoblast precursors are only differentiated at the early stage of the BM-MSC development, while adipogenesis is continuous across different stages.

### Osteoblast precursor induce angiogenesis during coupling with osteogenesis

Previous studies have reported that osteoblasts may regulate angiogenesis^39,40^, but this phenomenon has yet not been explored on the single-cell level. Interestingly, transcripts for some secreted factors associated with the vascular system (e.g., VCAN and ANGPTL4^41,42^ were highly expressed in osteoblast precursors, (**Supplementary Table 1: Sheet 3**). This result suggests that osteoblast precursors may induce angiogenesis concurrently with osteogenesis. In supporting this, the cluster marker genes of osteoblast precursors were enriched for not only osteogenesis related GO terms, but also for functional processes related to angiogenesis such as “regulation of vasculature development” and “positive regulation of angiogenesis” (**Fig. 1e, and 3a**). We further investigated the genes enriched for these biological processes and identified 32 genes regulating osteogenesis (e.g., COL1A1/A3, COL6A1/A3, VCAN, IGFBP3, etc.), 16 genes for angiogenesis (e.g., ADM, EGR1, NGFR, etc.), and 11 shared genes including MDK, JUNB, ENG, IGTB2, APOB, etc. (**Fig. 3b; Supplementary Table 3: Sheet 1**). Among these genes, some have a much higher expression level in osteoblast precursors compared with other cells.

Notably, we found that MDK, CD105, and ADAMTS9 were highly expressed and frequently enriched in multiple functional terms related to osteogenesis and angiogenesis (**Extended Data Fig. 3b**). It has been shown that MDK is positively associated with angiogenesis while inversely associated with osteogenesis^43,44^, potentially via MAPK and PI3K signaling^45^. High expression of CD105 has been shown to disrupt angiogenesis in tumor tissue, and CD105^-^ BM-MSCs are more prone to differentiate into adipocytes and osteocytes^11,46^. ADAMTS9 is expressed during ossification and also may regulate angiogenic signaling induced by VEGF^47,48^. Our results together with the previous evidence suggested that the co-regulation of osteogenesis and angiogenesis by osteoblast precursor is a complex network involving multiple genes whose regulation effect sometimes are in opposite directions.

The KEGG pathway analysis revealed that the osteogenesis and angiogenesis genes were enriched in the PI3K-Akt, MAPK, Rap1, AGE-RAGE, Relaxin, HIF-1, and TNF signaling pathways (**Fig. 3c**). The genes COL1A1, PGF, and JUN were highly expressed and were also enriched in multiple pathways, indicating that these genes may be essential in the cell signaling networks. We also found that PI3K-Akt signaling and osteogenesis share a large proportion of common genes, suggesting that this pathway may have a significant role in regulating the osteogenesis of BM-MSCs (**Fig. 3d**). On the other hand, the MAPK, PI3K-Akt, and Rap1 signaling pathways share comparable proportions of genes with angiogenesis (**Fig. 3d**). Furthermore, COL4A2, HGF, IGBT1, and ID1 were essential factors connecting the genetic network between the different pathways and biological processes (**Fig. 3e**). These results suggest that the osteogenesis and angiogenesis in osteoblast precursors may be mediated by multiple genes and pathways, particularly through PI3K-Akt and MAPK signaling.

### Myogenesis potential of CD56^**+**^ **chondrocyte precursors**

The DEGs in CD56^+^ chondrocyte precursors were enriched in GO terms related to both chondrogenesis (e.g., “cartilage development”, “chondrocyte differentiation”) and myogenesis (e.g., “muscle cell differentiation”, “myoblast differentiation”) (**Fig. 4a**). There were 46 DEGs enriched in terms related to chondrogenesis (e.g., IBSP, SPP1, A2M, IGTA10, etc.), 42 for myogenesis (e.g., ACTA2, ADARB1, CD9, VIM, etc.), and 13 shared genes for both processes (e.g., NPNT, MEF2C, ITGA8, TGFB1, etc.) (**Supplementary Table 3: Sheet 2**). Among the enriched genes, MEF2C and ITGA8 were highly expressed in CD56^+^ chondrocyte precursors and also related to multiple terms regarding chondrogenesis and myogenesis **(Fig. 4b; and Extended Data Fig. 3c**). MEF2C (myocyte enhancer factor 2C) is known to be essential for skeletal muscle development as well as attenuating MSC-derived cartilage hypertrophy in response to hypoxic conditioning^49,50^. ITGA8 (integrin subunit alpha 8) is a gene that modulates integrin activity to induce cartilage formation and protect against arthritis, while β1-integrin signaling enhances regeneration of myocytes^51,52^. Therefore, MEF2C and ITGA8 may be important drivers of the chondrogenesis and myogenesis potential for BM-MSCs.

**Fig. 4.**
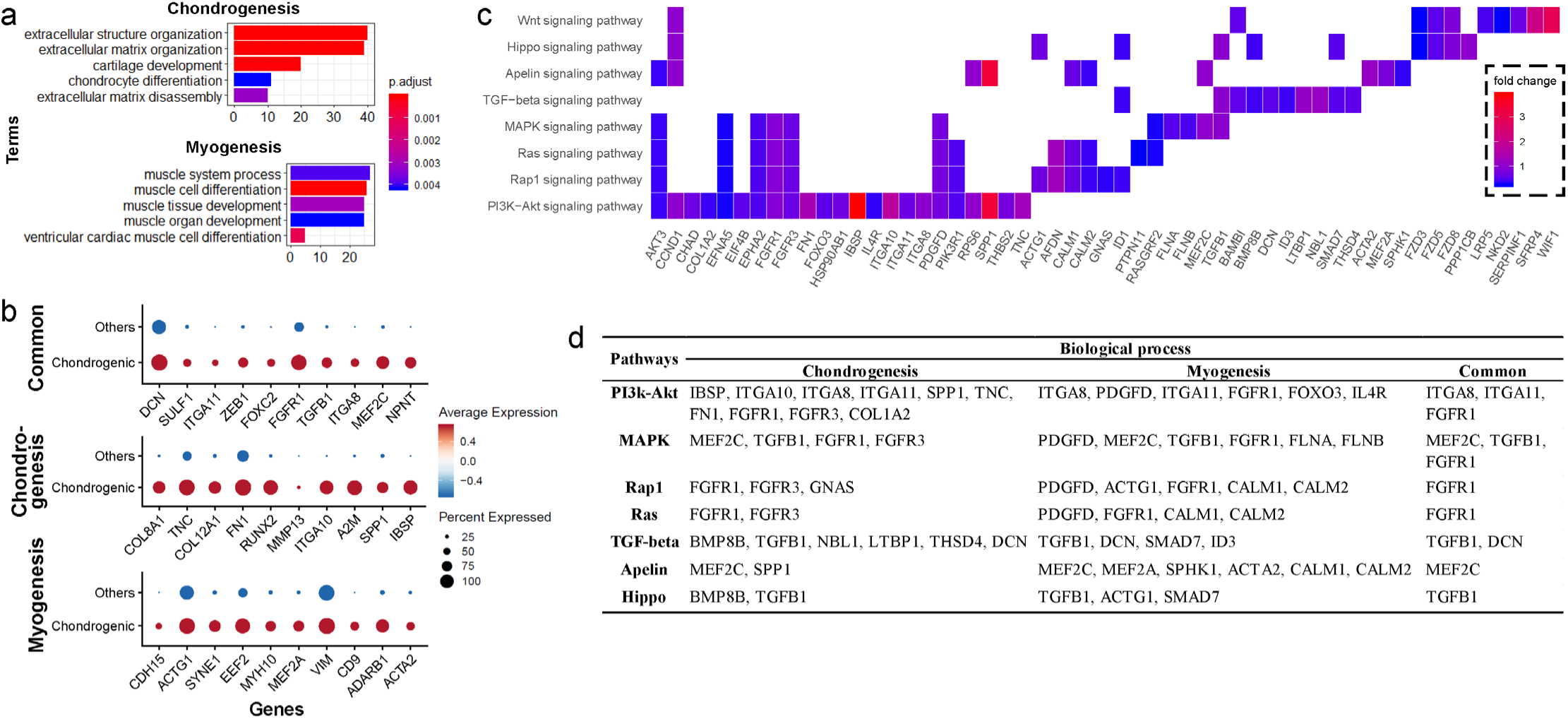
Functional analysis for CD56hi chondrocyte precursor. **a**, Enriched GO terms associated with chondrogenesis (top) and myogenesis (bottom) in chondrocyte precursor cells. Bar chart shows the number of enriched genes in each term. Color indicates the adjust *p* values. **b**, Differential expression of chondrogenesis and (or) myogenesis-related genes in chondrocyte precursors compared to the other cells. Dot plot shows the top 10 most-significant DEGs in each category (Middle: Chondrogenesis; Bottom: Myogenesis; Top: Common for both), where dot color indicates the relative expression levels between chondrocyte precursors and other cells (z-score) and the dot size shows the proportion of cells expressing the indicated genes. **c**, Gene expression pattern in enriched pathways. Squares show enriched DEGs in the corresponding terms (rows). Color indicates the expression value of the DEGs (average logFC). **d**, Table of genes in biological processes and pathways.

Based on the KEGG pathway analysis, we determined that DEGs in the chondrocyte precursors were enriched in the PI3K-Akt, MAPK, Ras, Rap1, TGF-beta, Apelin, and Hippo signaling pathways (**Fig. 4c**). TGF-beta signaling shared the largest number of genes with chondrogenesis, while the genes enriched in Apelin and Ras/Rap1 signaling overlapped most with myogenesis (**Extended Data Fig. 3d**). By investigating the overlapping genes between biological processes and signaling pathways, we found that FGFR1 and TGFB1 may be crucial genes connecting multiple pathways to both chondrogenesis and myogenesis (**Fig. 4d**). Thus, the CD56^+^ chondrocyte precursor of BM-MSC subpopulation is capable of both chondrogenesis and myogenesis, and these processes may be regulated by the TGF-beta, Apelin, and Ras/Rap1 signaling pathways.

### Transcriptional difference between human and mice BM-MSCs at single-cell level

To investigate the difference of transcriptional profiles between BM-MSCs originated from human and mice (hBM-MSCs, mBM-MSCs, respectively), we integrated our single cell human transcriptome data from two previous scRNA-seq studies on bone marrow components^7,19^. Potential batch effects among different studies were reduced by a canonical correlation analysis (CCA) (**see methods**)^53,54^, and different datasets had a high correlation (**Fig. 5a-c**), suggesting that after the CCA integration, the batch effects between different studies were relatively small and were, therefore, less likely to introduce notable bias into the downstream analysis.

**Fig. 5.**
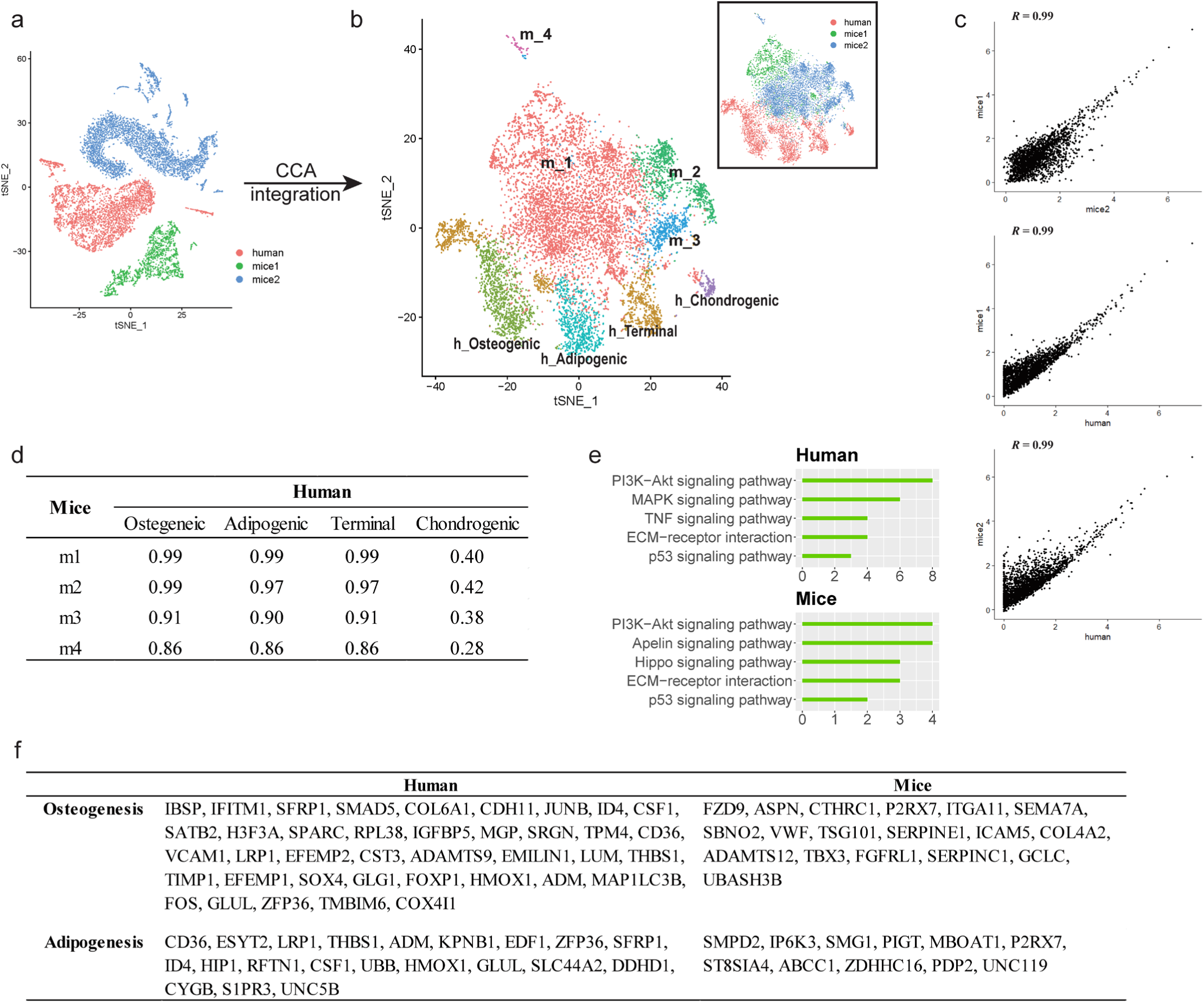
Integrated cross-species analysis between human and mouse BM-MSCs. **a-b**, t-SNE visualization of human and mouse BM-MNCs before (**a**) and after (**b**) CCA integration. The labeled texts indicate the datasets (**a**, and box in **b**) or subpopulations identified by clustering analysis (main plot in **b**). Human (h): data from this study; mice1 (m): data from Tikhonova et al.^7^; mice2 (m): data from Baryawno et al.^19^. Clusters: Osteogenic, Chondrogenic, adipogenic and terminal in human; m1-m4 in mice. **c**. Correlations of gene expression among different BM-MSC datasets after CCA integration. Each dot represents an individual gene. The average gene expression level (logFC)s are plotted for each subject. Correlations were measured by Pearson correlation coefficients (*R*) (*p* < 0.01). **d**, Correlations of gene expression between different subsets of human and mouse BM-MSCs identified by clustering analysis (Osteogenic, chondrogenic, adipogenic and terminal in human; m1-m4 in mice). Values in the table represent the Pearson correlation coefficients (*R, p* < 0.01). **e**, Enriched signal pathways (KEGG terms) of human (top) and mice (bottom) BM-MSCs. Bar chart shows the number of enriched genes in each term. **f**, Table comparing enriched genes in osteogenesis- and adipogenesis- related terms between human and mice BM-MSCs.

To test whether heterogeneity exists between human and mice BM-MSCs, integrated cross-species data was analyzed by an unbiased clustering. hBM-MSCs and mBM-MSCs were separated into different clusters (Osteogenic, Chondrogenic, adipogenic and terminal in human; m1-m4 in mice) (**Fig. 5b**). The clustering suggests that even though the overall data have a good correlation based on average gene expression, there are still systematic difference of transcriptome between hBM-MSCs and mBM-MSCs at the single-cell level. There was a strong correlation between the average gene expression of subtypes in hBM-MSCs and mBM-MSCs except for human chondrogenic BM-MSCs (**Fig. 5d**). This observation suggests that the overall gene expression pattern and differentiation trajectory of hBM-MSC-derived chondrocyte precursors is less similar with those in the mBM-MSCs, when compared to other hBM-MSC subpopulations.

Human and mice BM-MSCs often present different cell surface markers^3^. Consistent with this result, by comparing the DEGs between hBM-MSCs and mBM-MSCs (**Extended Data Fig. 4, Extended Data Fig. Table 1: Sheet 4**), we revealed several CD markers with distinct expression patterns between human and mice BM-MSCs. For instance, CD317, CD36, and CD63 were highly expressed in hBM-MSCs, but not in mBM-MSCs; and *vice versa* for CD148, CD108, and CD20 (**Extended Data Fig. 4d**). Next, we examined the difference DEGs associated with biological processes or signal pathways (**Supplementary Table 2: Sheets 3-4**). PI3K-Akt, p53, and ECM signaling were highly enriched in both human and mice BM-MSCs. MAPK and TNF signaling were highly enriched only in hBM-MSCs, but not in mBM-MSCs, *vice versa* for Apelin and Hippo signaling in mBM-MSCs (**Fig. 5e**).

Interestingly, when comparing the DEGs enriched in the osteogenic and adipogenic GO terms between hBM-MSCs and mBM-MSCs, no shared gene between human and mice BM-MSCs was observed (**Fig. 5f**), suggesting that though hBM-MSCs and mBM-MSCs shared the same biological processes, the regulating genes and pathways or their relative significance during the processes could be different. Overall, these results clearly demonstrated that there are considerable systemic differences in the transcriptional profiles between hBM-MSCs and mBM-MSCs at the single-cell level, which may provide novel insights into the biological basis underlying the fundamental differences in phenotypes and responses to various conditions in humans and mice^55,56^.

## Discussion

While a growing body of evidence indicates that BM-MSCs have a central role in bone health, the underlying subtypes of BM-MSCs, especially *in vivo* in humans, remains largely unknown due to its heterogeneous characteristics. In the present study, we applied scRNA-seq analysis on freshly isolated human BM-MSCs and their niche hematopoietic cells. The use of freshly isolated human cells is a major advantage of this study since any form of extra *in vitro* operations (e.g., freezing, culturing) could potentially alter the true transcription pattern^22^ and thus leading to biased cell clustering/identification. In addition, our results along with previous evidence have highlighted that transcription profiles vary largely between humans and mice^23^.

Several studies have applied scRNA-seq in bone marrow stroma components or MSCs derived from various origin (e.g., bone marrow, adipocytes, umbilical cord). For instance, Tikhonova et al.^7^ and Baryawno et al.^19^ independently performed scRNA-seq in bone marrow stroma components (including BM-MSCs, vasculature, osteoblastic cells, etc.). Similar to their results, we also identified subtypes corresponding to multiple trajectories in BM-MSCs. Chan et al.^15,16^, on the other hand, identified skeletal stem cells in humans and mice. They also demonstrated a Lin^-^PDPN^-^CD146^+^ osteogenic subsets that only give rise to osteoblasts/osteocytes^15^. Similarly, we found that CD146 was differentially expressed in osteogenic subset of BM-MSCs. Some studies also performed scRNA-seq on cultured human MSCs derived from various origins^20,21,57^, but none of these studies focused on subtype identification. Compared with these studies which focused on mice cells or *in vitro* cultured human cells, our results thus greatly expand the understanding of *in vivo* human BM-MSCs by presenting an unbiased transcriptional profile of distinct subpopulations including osteoblast, chondrocyte and adipocyte precursors as well as, other components of the human BM-MSC cell population *in vivo*.

Although the use of freshly isolated cells for scRNA-seq may preserve to the largest extent the accuracy of the transcriptomic profile, this approach also limits the total number of collected cells. Therefore, we used a single marker – CD271 – for positive sorting, instead of combining with CD45- negative selection which, would generate even less yield. Based on the scRNA-seq gene expression profiles, we demonstrated that the CD271^+^ BM-MNCs represent a heterogeneous cell population which may be subdivided into BM-MSCs along with various hematopoietic cells which contribute to the formation of niche components. Our finding suggests that the BM-MSC isolation protocol based solely on positive selection is not ideal since the isolated cells consist of various cell types. Instead, positive selection combined with negative selection using CD45 or lineage markers (LIN) should be considered if the major purpose is to isolate BM-MSC with the higher purity^5,58^.

Since BM-MSCs are heterogeneous for several existing cell markers^7,9^, it is necessary to search for novel BM-MSC-specific cell markers (specifically and uniformly expressed in the major BM-MSC population). By comparing the expression pattern between BM-MSCs and other niche hematopoietic cells, we confirmed the expression of classic cell markers including CD271, LEPR, CD105, CD90 at the single-cell level. Notably, we found that LEPR had the highest expression level and was specific to the BM-MSC population, which is consistent with the results from mouse models^24^. We also detected some additional specifically expressed CD markers (e.g., CD167b, CD91, CD130, CD118) in BM-MSCs, which may potentially serve as novel surface markers for BM-MSC enrichment/purification.

A systematic analysis of the BM-MSC transcriptional profiles revealed distinct subpopulations corresponding to osteogenic, chondrogenic, and adipogenic differentiation, as well as terminal-stage cells in the quiescent state. Further examination into the relationships between the highly expressed genes, biological processes, and signaling pathways in each subpopulation suggested that osteoblast precursors may have the capacity to induce vasculature development, and the chondrocyte precursors may have myogenic potential. Normally, the coupling of osteogenesis and angiogenesis is in the same regulation direction, i.e., vascular development will promote bone formation and *vice versa*^59^. However, some recent studies have already shown that in some cases the regulation effect of these two biological processes could be opposite. For instance, even though VEGF stimulates vascularization, high amount of VEGF could impair bone formation^60^. Similar patterns were found in BM-MSCs in this study where osteoblast precursors express CD105 and MDK, whose regulation effect on osteogenesis and angiogenesis may be opposite, suggesting that the coupling of osteogenesis and angiogenesis is a complex regulation network where both positive and negative feedback may be included.

The scRNA-seq profiles of the BM-MSCs also revealed a continuum of dynamic gene expression pattern, indicating that osteogenesis only occurs at early stages of BM-MSC development while the adipogenic and quiescent cell take a dominate place in the terminal stages (**Fig. 2b**). These findings suggested that the aging of BM-MSCs may be an important factor in the balance between the osteogenic and adipogenic differentiation.

While the overall data did not show a significant batch effect, the transcription pattern of the BM-MSCs varied largely between subjects (**Extended Data Fig. 2d**). Counterintuitively, the osteogenic subpopulation was mostly derived of cells from the low bone mineral density (BMD) osteoporotic female subject, whereas the adipogenic subpopulation was primarily composed of cells from the osteoarthritic male subject with normal BMD (**Fig. 1c**). We hypothesize that this may be at least partially explained by the gender and age difference between the two subjects. Females are expected to suffer from osteoporosis earlier in life compared to males^61^. A primary etiology of postmenopausal osteoporosis is an estrogen-deficiency induced elevated number and activity of osteoclasts, which are responsible for resorption of bone tissue, while the number of osteoblasts remains relatively constant or may even increase to compensate for bone loss, when comparing to the age-matched controls^62,63^. Though in male the number and the activity of osteoclast remains the same, with increasing age, the production and activity of osteoblasts is dramatically reduced^2^. The osteoporosis subject in this study is a 67-year-old postmenopausal female, while the osteoarthritis subject is an 85-year-old male. Therefore, the large discrepancy in age and the gender difference rather than disease status may largely underlie the observed difference in transcriptomic profiles of the BM-MSCs between the two subjects. However, further investigations are needed to determine whether and/or how such differences are related to the disease status (osteoporosis vs. osteoarthritis) or other factors (e.g., age, gender, lifestyle, medical/medication history).

Despite providing a detailed characterization of human BM-MSCs at single-cell resolution, the full trajectory of the osteoblastic lineage cells, as well as their balance and interaction with the osteoclastic lineage remains elusive. In our future studies, by combining scRNA-seq with scATAC-seq – a powerful tool to evaluate chromatin accessibility at the single-cell level, we aim to unveil the complexity of osteoblastic-osteoclastic lineage interactions and gene expression regulations with/between the two lineages. In the meantime, deconvoluting the heterogeneity of the BM-MSCs *in vivo* in humans represents an important, first, and necessary advancement towards improving our understanding of bone physiological processes.

## Methods

### Study population

The clinical study was approved by the Medical Ethics Committee of Central South University, and written informed consents were obtained from each participant. The study population consists of two Chinese subjects who underwent hip replacement surgery at the Xiangya Hospital of Central South University in 2019, including one 67 year old female with osteoporosis (BMD T-score: -3.3 at lumbar vertebrae, -3.7 at left hip joint, list specific bone areas for DXA measurements) and one 84 year old male with osteoarthritis and normal BMD (BMD T-score: -0.9 at lumbar vertebrae, 2.7 at left hip joint). Study participants were screened prior to surgery answering a detailed questionnaire, completing a medical history, and a physical examination. Subjects were excluded from the study if they had preexisting chronic conditions which may influence bone metabolism including diabetes mellitus, renal failure, liver failure, hematologic diseases, disorders of the thyroid/parathyroid, malabsorption syndrome, malignant tumors, and previous pathologic fractures^64^. During hip replacement surgery, physicians collected the bone marrow from the femoral shafts from each subject and transferred the samples to our laboratory immediately following the procedure. The samples were stored at 4°C and processed within 24 hours after collection.

### BMD measurement

BMD (g/cm^2^) at the lumbar spine (L1–L4) and the left hip was measured with a duel energy x-ray absorptiometry (DXA) fan-beam bone densitometer (Hologic QDR 4500A, Hologic, Inc., Bedford, MA, USA). According to the World Health Organization definition^65^ and the BMD reference established for Chinese populations^66^, subjects with a BMD of 2.5 SDs lower than the peak mean of the same gender (T-score ≤ -2.5) were determined to be osteoporotic, while subjects with -2.5 < T-score < -1 are classified as having osteopenia and subjects with T-score > -1.0 are considered healthy.

### Bone marrow cell dissociation

Bone marrow derived mononuclear cells (BM-MNCs) were extracted from the marrow cavity of femoral shafts using a widely applied dissociation protocol^5,6^. Briefly, the bone marrow was attenuated with PBS (1:2) and mixed gently. The mixture was then equally layered onto equal volume of Ficoll (GE health care, Chicago, IL, USA), and the buffy coat was isolated by centrifugation (440g, 35 min, 4°C). The separated buffy coat was transferred into a new 15 ml centrifuge tube and washed with PBS. After discarding the supernatant, red blood cells were lysed with RBC Lysis Buffer (Thermo Fisher, Waltham, MA, USA). After washing twice with PBS, the remaining MNCs were further purified with CD271 magnetic MicroBeads (Miltenyi Biotec, Bergisch Gladbach, Germany) for positive selection^6^.

### Positive selection of CD271^**+**^ **BM**-**MNC**

BM-MNCs were incubated for 10 min at 4–8°C with monoclonal antibody (mAb) against CD271. After washing, the cells were incubated with anti-IgG1 immunomagnetic beads for 15 min at 4°C. The cell suspension was placed on a column in a cell separator (Miltenyi Biotec), and the positive fraction was subjected to a second separation step. The cells were then counted and assessed for viability with a Countstar® Rigel S3 fluorescence cell analyzer (ALIT Life Science Co., Ltd, Shanghai, China).

### Cell capture and cDNA synthesis

After cell isolation, we applied the Chromium single cell gene expression platform (10x Genomics, Pleasanton, CA, USA) for the scRNA-seq experiments. Cell suspensions were loaded on a Chromium Single Cell Controller (10x Genomics) to generate single-cell gel beads in emulsion (GEMs) by using Single Cell 3’ Library and Gel Bead Kit V3 (10x Genomics, Cat# 1000092) and Chromium Single Cell A Chip Kit (10x Genomics, Cat#120236) according to the manufacturer’s protocol. Briefly, single cells were suspended in 0.04% BSA–PBS. Cells were added to each channel, captured cells were lysed, and the released RNA were barcoded through reverse transcription in individual GEMs^67^. GEMs were reverse transcribed in a C1000 Touch Thermal Cycler (Bio Rad, Hercules, CA, USA) programmed at 53°C for 45 min, 85°C for 5 min, and held at 4°C. After reverse transcription, single-cell droplets were broken, and the single-strand cDNA was isolated and cleaned with Cleanup Mix containing DynaBeads (Thermo Fisher Scientific). The cDNA was generated and amplified, and quality was assessed using the Agilent 4200.

### Single cell RNA-Seq library preparation

Single-cell RNA-seq libraries were prepared using Single Cell 3’ Library Gel Bead Kit V3 following the manufacturer’s guide (https://support.10xgenomics.com/single-cell-gene-expression/library-prep/doc/user-guide-chromium-single-cell-3-reagent-kits-user-guide-v3-chemistry). Single Cell 3’ Libraries contain the P5 and P7 primers used in Illumina bridge amplification PCR. The 10x Barcode and Read 1 (primer site for sequencing read 1) were added to the molecules during the GEM-RT incubation. The P5 primer, Read 2 (primer site for sequencing read 2), Sample Index and P7 primer were added during library construction. The protocol is designed to support library construction from a wide range of cDNA amplification yields spanning from 2 ng to >2 μg without modification. Finally, sequencing was performed on an Illumina Novaseq6000 with a sequencing depth of at least 100,000 reads per cell for a 150bp paired end (PE150) run.

### Pre-processing of scRNA-seq data

Raw FASTQ files were mapped to the Reference genome (GRCh38/hg38) using Cell Ranger 3.0 (https://support.10xgenomics.com/single-cell-gene-expression/software/pipelines/latest/what-is-cell-ranger). To create Cell Ranger-compatible reference genomes, the references were rebuilt according to instructions from 10x Genomics (https://www.10xgenomics.com), which performs alignment, filtering, barcode counting and UMI counting. Following alignment, digital gene expression (DGE) matrices were generated for each sample and for all samples. Merged 10x Genomics DGE files were generated using the aggregation function of the Cell Ranger pipeline. All the cells in different batches were merged and normalized by equalizing the read depth among libraries. Only confidently mapped, non-PCR duplicates with valid barcodes and unique molecular identifiers were used to generate the gene-barcode matrix (**Extended Data Fig. 1a**,**b**). For further quality control, we excluded cells that had fewer than 150 detected genes. We then calculated the distribution of genes detected per cell and removed any cells in the top 2% quantile. We also removed cells where > 20% of the transcripts were attributed to mitochondrial genes (**Extended Data Fig. 1c**,**d**). After removing disqualified cells from the dataset, the data was normalized by the total expression, multiplied by a scale factor of 10,000, and log transformed.

### Dimensionality reduction and data visualization

To visualize the data, we first calculated the ratio of binned variance to mean expression for each gene and selected the top 2,000 most variable genes. Next, we performed principal component analysis (PCA) and reduced the data to the top 20 PCs. Finally, we performed non-linear dimensionality reduction for the dataset to project the cells in 2D space based on gene expression data of the highly variable genes using t-SNE^68^.

### Clustering and differential gene expression analysis

We performed a graph-based clustering of the previously identified PCs using the Louvain Method^69^, and the clusters were visualized on a 2D map produced with t-SNE. For each cluster, we used the Wilcoxon rank-sum test to identify significantly differentially expressed genes (DEGs) when compared to the remaining clusters (multiple hypothesis testing was adjusted by Bonferroni correction, adjusted *p* value < 0.05 was regarded as significant, paired tests when indicated). To visualize how well the cluster-specific DEGs (marker genes) defined each cluster, we constructed the violin plot, feature plot (tSNE plot colored by expression level of indicated genes), and heatmap (top 10 genes with highest average log-transformed fold change – logFC) using the *Seurat* R packages^70,71^.

### Pathway enrichment analysis and trajectories analysis

To investigate the biological processes and signaling pathways associated with each cluster (subtype), we performed GO and KEGG enrichment analysis on the identified cluster-specific DEGs by using the *clusterProfiler* R package^72^. To visualize the results, we used the *ComplexHeatmap* and *GOplot* R packages. We then applied *Monocle* for trajectory inference and pseudotime analysis^73,74^. The principle of these analyses is to determine the pattern of the dynamic process experienced by the cell population and to order the cells along their developmental trajectory based on differences in the expression profiles of highly variable genes.

### Cross-species scRNA-seq data integration

Two previous scRNA-seq data of mBM-MSCs were acquired from GEO database under the accession numbers of GSE128423 and GSE108892, respectively^7,19^. After acquiring expression matrix, cells expressing LEPR was isolated as LEPR^+^ mBM-MSC subset. Then we applied a canonical correlation analysis (CCA) using Seurat alignment method to integrate scRNA-seq data of hBM-MSCs and mBM- MSCs^70,71^. The CCA methods is to find the linear combinations of features, and then identifies shared correlation structures across different datasets. First, we identified variable genes and controlled for the strong relationship between variability and average expression for each dataset. Then, we selected the top 2,000 genes with the highest dispersion shared between each dataset and ran the CCA to determine the common sources of variation. Finally, we aligned the subspaces based on the first 30 canonical correlation vectors, which result in a new dimensional reduction that was used for further analysis^7^. The batch effect was then tested using a coefficient analysis of average gene expression between each of the datasets.

## Acknowledgement

This research was benefited by grants from the National Institutes of Health (R01AR069055, U19AG055373, P20GM109036, R01AG061917), the Franklin D. Dickson/Missouri Endowment, and the Edward G. Schlieder Endowment and the Drs. W. C. Tsai and P. T. Kung Professorship in Biostatistics from Tulane University, Special Funding for the Construction of Innovative Provinces in Hunan (Grant No. 2019SK2141), China Oceanwide Holding Group Project Fund (Contract No.143010100), National Natural Science Foundation of China (Grant No. 81902277) and Central South University (Grant No. 2018zzts886). We are thankful to our cooperators Dr. Wei Liu, Dr. Xiaoshan Tian, and Mr. Qing-Zhong Hua, who provided expertise that greatly assisted the research.

## Author contributions

ZW wrote the main manuscript text and conducted major analysis; CL, JX, XY, and YL collected the human sample and corresponding clinical information; XL, JY, YH, and HZ performed the experiments; JG, HS, and MRS did language proofreading; YG, LJT, SYT prepared supplementary information and validated the results; the study was conceived, designed, initiated, directed and supervised by HS, HMX, and HWD. All authors participated in the discussions of the project and reviewed and/or revised the manuscript.

## Data availability

Two previous scRNA-seq data of murine BM-MSCs used in this study are available in the GEO database with accession numbers GSE128423 and GSE108892. The scRNA-seq data of CD271^+^ bone marrow mononuclear cells from two human sample can be accessed with accession number under GSE147287, which is embargoed until publication of this study.

## Extended Data Figures

**Extended Data Fig. 1.**
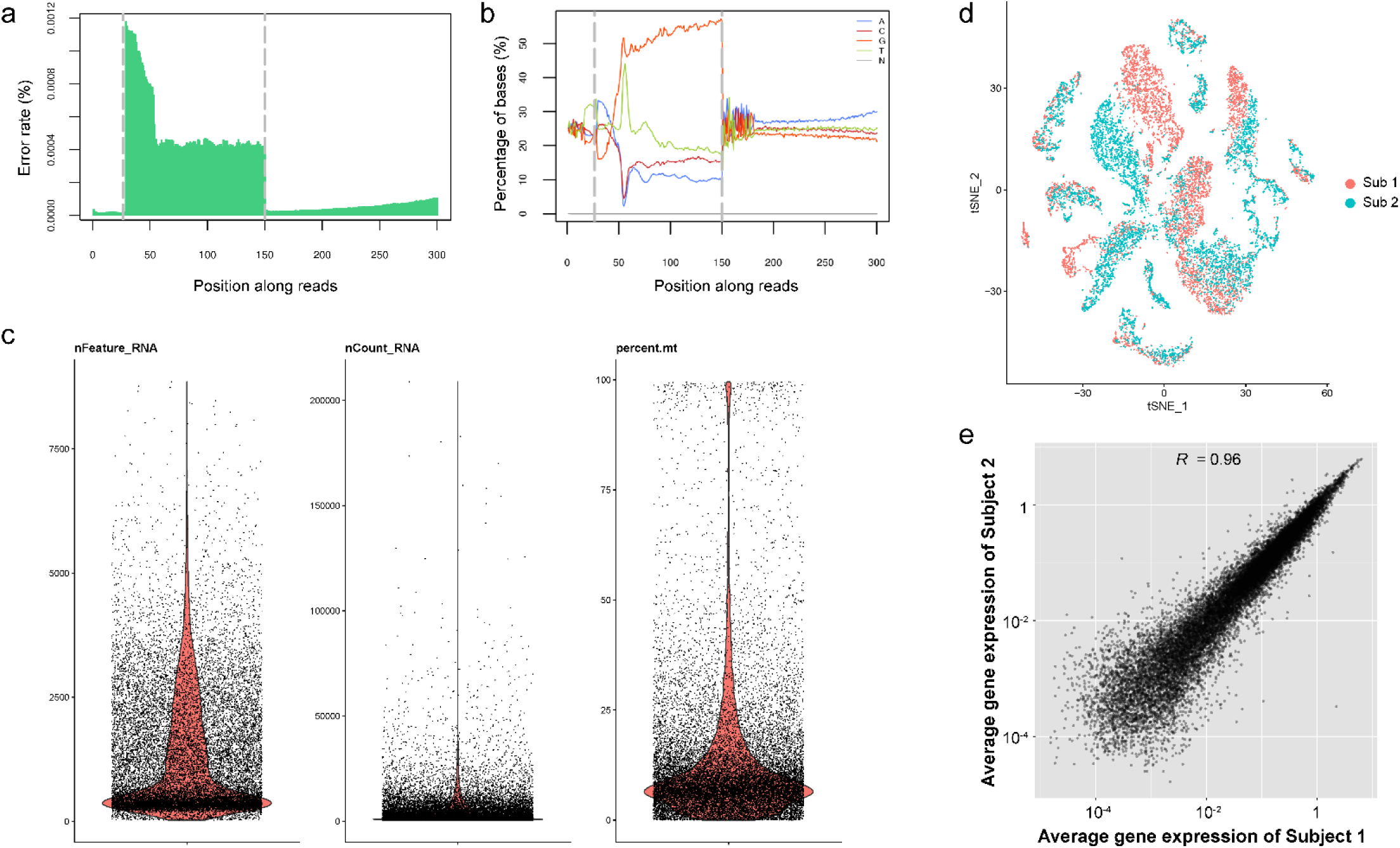
Quality control for scRNA-seq data. **a**, Average error rate distribution of base pair. x-axis represents position of base pair along reads while y-axis represents the error rate. Dashed line at 150 bp separates Read1 and Read2 (barcode and RNA sequence, representative). The average error rate of Read2 (RNA sequence) was less than 1×10^−4^. **b**, Base content distribution. x-axis represents position of base pair along reads while y-axis represents the proportion of different nucleotides (A, T, G, C, N stands for uncertain). Dashed line at 150 bp separates Read1 and Read2. All the bases were determined to A, T, G, or C with no undetermined bases, and difference between A and T, or G and C is less than 10% in any position. Combined with (A), it suggests a good quality of raw sequencing data. **c**, Distribution of the number of detected genes (features), RNA molecules, and proportion of mitochondrial genes (left to right). Mean Reads per Cell: 5,458; Mean Genes per Cell: 1,339. In less than 20% of cells that proportion of mitochondrial genes is over 20%, therefore in the majority of the cells, most of the detected reads and genes are from the genome. **d**, t-SNE dimension reduction, colored by different subjects (sub1, osteoporosis, red, n = 7,498; sub2, osteopenia, blue n = 6,996). **e**, Correlation of gene expression between two subjects. Each dot represents an individual gene. Axis measure the average gene expression level in the indicated subject (axis is log-scaled). Correlation was tested by Pearson correlation coefficient (*R* = 0.96, *p* < 0.01).

**Extended Data Fig. 2.**
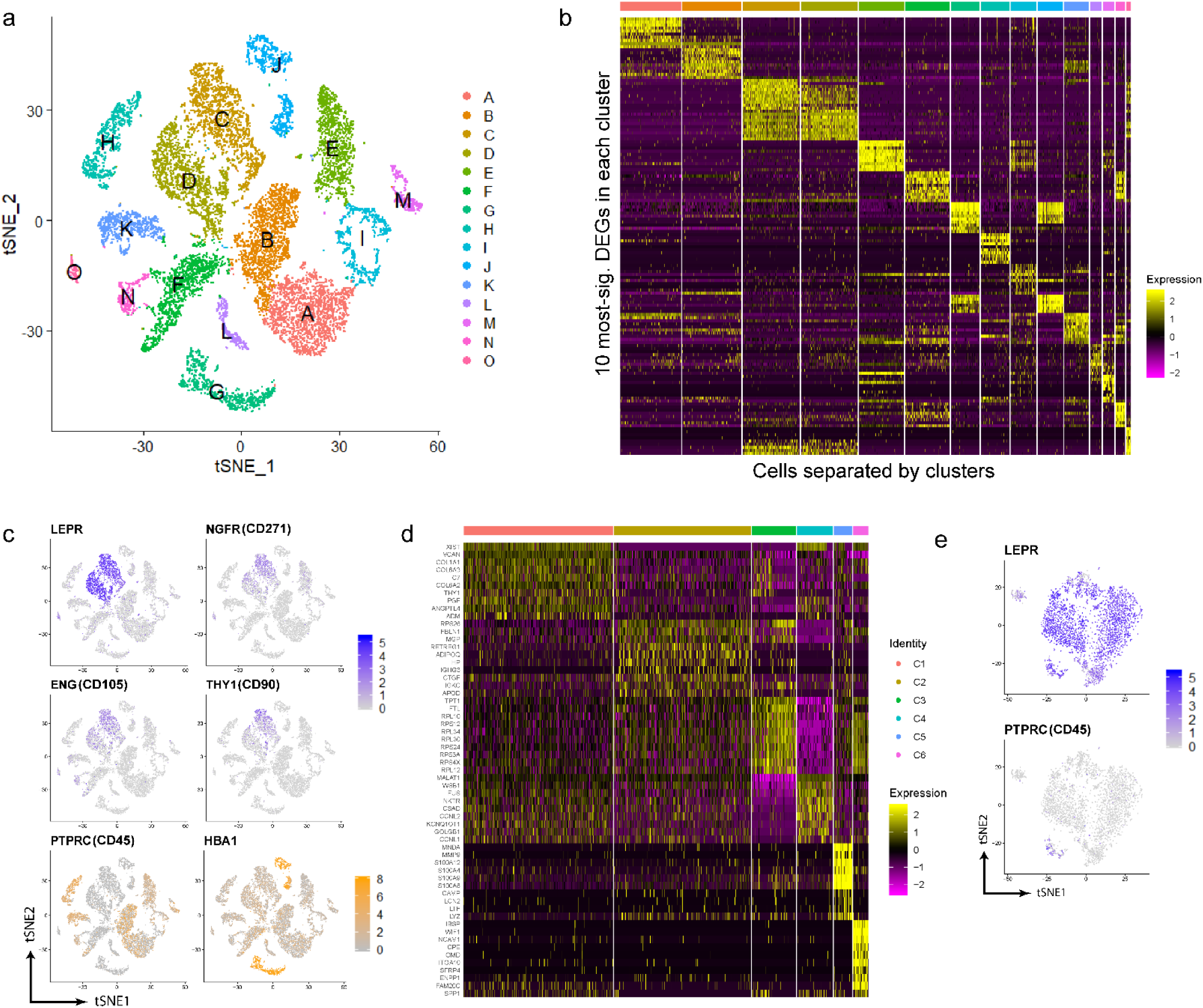
Cluster and marker gene identification for scRNA-seq data. **a**, t-SNE dimension reduction, colored by different clusters. **b**, Gene expression profile of CD271^+^ BM-MNCs, based on the relative gene expression level of top 10 most-significant markers for each cluster. **c**, Gene expression of known BM-MSC markers, embedded on t- SNE dimension reduction map, and colored by gene expression level. **d**, Gene signature of BM-MSCs, based on the relative gene expression level of top 10 most-significant DEGs for each cluster (z-score). **e**, Gene expression of LEPR and PTPRC (CD45) in BM-MSCs, embedded on t-SNE dimension reduction map, and colored by gene expression level (logFC).

**Extended Data Fig. 3.**
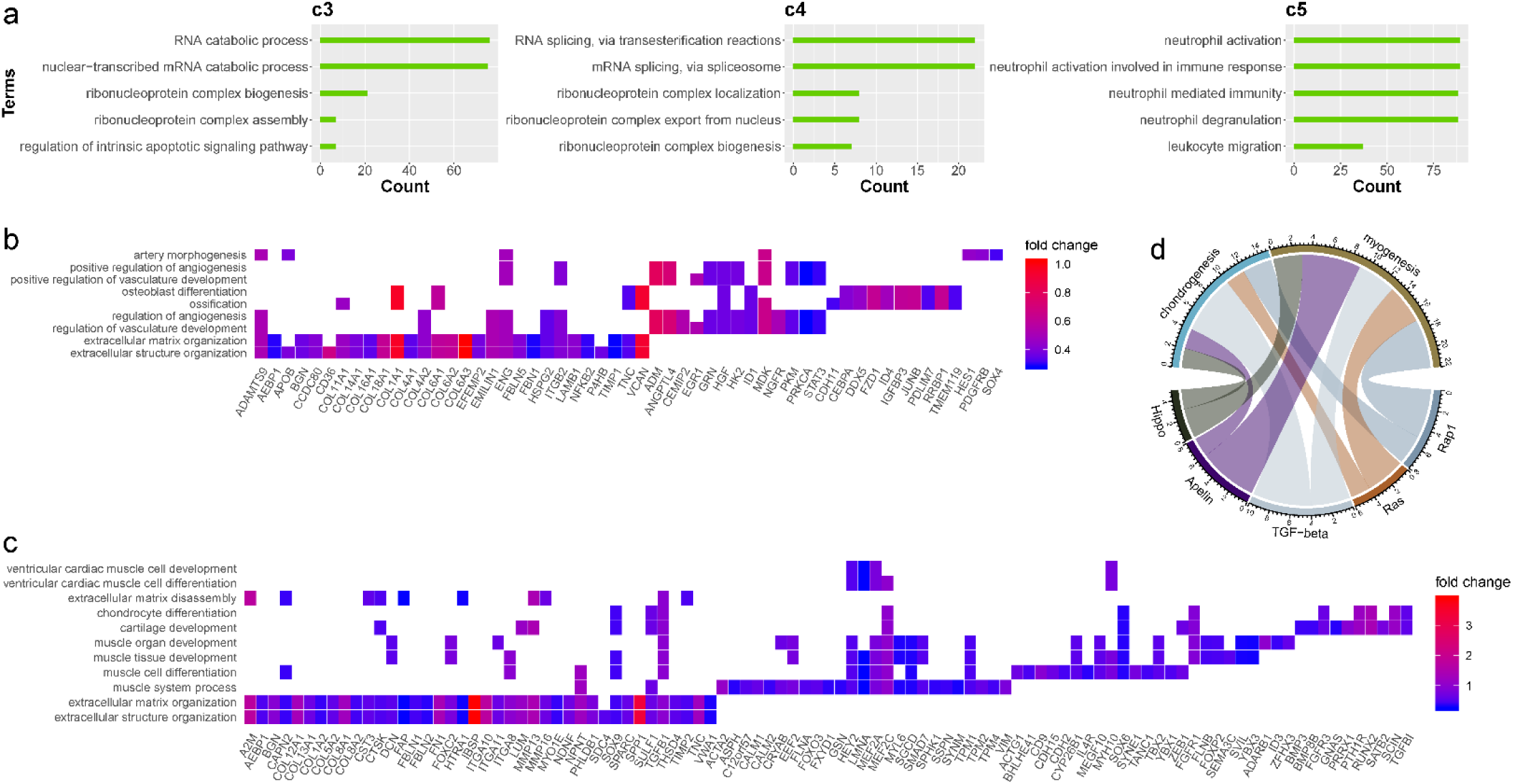
GO enrichment for BM-MSC. **a**, Enriched GO terms in cluster 3 to 5 (left, middle, right, respectively). Bar chart shows the number of enriched genes in each term. **b-c**, Gene expression pattern in enriched pathways for osteoblast (**b**) and chondrocyte (**c**) precursors. Squares showing enrich DEGs in the corresponding terms (rows). Color indicating the gene expression level (average logFC). **d**, Common genes shared between biological processes and pathways. Width of curves connecting different terms indicating the relative proportion of the shared genes.

**Extended Data Fig. 4.**
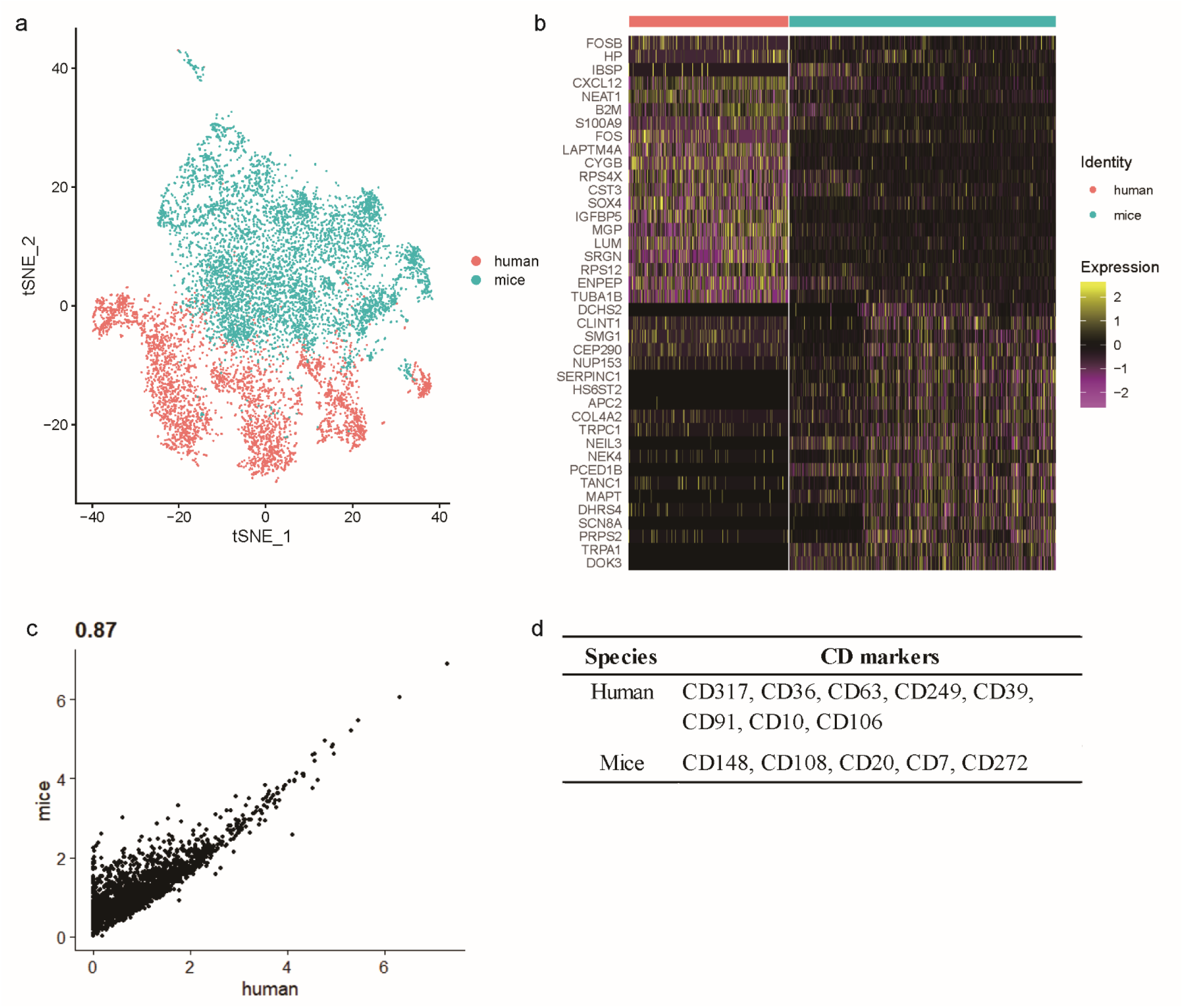
Comparison of gene expression profile between hBM-MSCs and mBM-MSCs. **a**, t-SNE visualization of human and mouse BM-MNCs, colored by species. **b**, Gene signature of human and mice BM-MSCs, based on the relative gene expression level of top 20 most-significant DEGs for each species (z-score). **c**, Correlation of gene expression between human and mice. Each dot represents an individual gene. Axis measure the average gene expression level (logFC) in the indicated species. Correlation was tested by Pearson correlation coefficient (*R* = 0.87, *p* < 0.01). **d**, Comparison of differentially expressed CD markers between human and mice BM-MSCs.

