## Supplementary information for "Single-cell RNA sequencing deconvolutes the *in vivo* heterogeneity of human bone marrow-derived mesenchymal stem cells"

**Supplementary** **Table 1**. Differential gene expression analysis. Sheet 1: Cluster-specific DEGs of each cluster in CD271^+^ BM-MSCs. Sheet 2: DEGs in LEPR^hi^CD45^low^ BM-MSCs when comparing with other CD45^hi^ BM-MNCs. Sheet 3: Cluster-specific DEGs for BM-MSC clusters. Sheet 4: Relative gene expression between hBM-MSCs and mBM-MSCs. Average log-transformed fold change (logFC) was calculated on averaging the expression value of all single cells in each cluster against other clusters using Wilcoxon rank-sum test.

**Supplementary Table 2.** Enriched GO and KEGG terms for BM-MSCs. Sheets 1-2: Enriched GO and KEGG terms of cluster-specific DEGs six identified clusters in BM-MSCs, respectively. Sheets 3-4, Enriched GO and KEGG terms of DEGs between hBM-MSCs and mBM-MSCs, respectively.

**Supplementary Table 3.** Expression pattern of enriched genes in related biological processes. Sheets 1-2: Enriched genes in osteoblast and chondrocyte precursors, respectively.
